# Isoform-specific single-cell perturb-seq reveals distinct functions of alternative promoters in drug response

**DOI:** 10.1101/2025.07.14.664827

**Authors:** Helen E. King, Savannah O’Connell, Daisy Kavanagh, Sofia Mason, Cerys McCool, Javier Fernandez-Chamorro, Christine Chaffer, Susan J Clark, Helaine Graziele S. Vieira, Timothy Sterne-Weiler, Robert J Weatheritt

**Author notes:** Senior Authors (Helaine Graziele S. Vieira, Timothy Sterne-Weiler, Robert J Weatheritt).

## Abstract

CRISPR interference (CRISPRi) screens have emerged as powerful tools for dissecting gene function, yet their application to genes with multiple promoters, which comprise over 60% of human genes, remains poorly understood. Here, we demonstrate that CRISPR-dCas9-based screens exhibit widespread promoter specificity, with untargeted promoters often showing compensatory upregulation to maintain gene expression. Leveraging this selective targeting of individual promoters within the same gene, we developed isoform-specific single-cell Perturb-Seq to systematically analyse alternative promoter function. Our analysis revealed that alternative promoters in 48.3% of targeted genes drive distinct transcriptional programs. This suggests that promoter selection represents a fundamental mechanism for generating cellular diversity rather than mere transcriptional redundancy. In breast cancer models, this promoter-specific targeting revealed differential effects on drug sensitivity, where distinct estrogen receptor (ESR1) promoters showed opposing influences on tamoxifen response and patient survival. These findings demonstrate the necessity of promoter-level analysis in functional genomics and suggest new strategies for therapeutic intervention through promoter-specific targeting.

## Introduction

CRISPR interference (CRISPRi) has revolutionised functional genomics by enabling the precise control of gene expression through sequence-specific targeting of promoter regions(1). This technology has uncovered complex cellular networks and mechanisms, ranging from basic biological processes(2-4) to therapeutic applications(5-8). The scalability of these screens, combined with their ability to tune gene expression rather than completely ablate it, has opened new avenues for understanding complex biological processes and developing more effective therapeutic strategies(1,4). However, a critical limitation has emerged: current approaches predominantly target a single promoter per gene(1-4,6,7,9), potentially missing crucial regulatory dynamics in the over 60% of human genes that utilize multiple promoters(10,11).

Alternative promoters serve as molecular switches, enabling genes to generate distinct transcript isoforms in response to cellular demands and environmental signals. Each promoter region integrates a wide array of regulatory signals, including from distal enhancers, chromatin modifications, and transcription factor binding, to control gene expression in di\erent cellular contexts(12,13). Growing evidence suggests these alternative promoters play crucial roles in development, disease progression, and therapeutic response, yet their systematic functional analysis has remained technically challenging.

The CRISPRi machinery typically influences transcription within approximately 1000 nucleotides of the guide RNA binding site, suggesting that distally separated alternative promoters might be independently targetable(14). This spatial specificity, combined with the prevalence of alternative promoters separated by more than 1000 nucleotides, presents both a challenge and an opportunity: while current CRISPRi libraries may miss critical regulatory elements, the technology could potentially enable the systematic analysis of alternative promoter function.

Here, we leverage the spatial specificity of CRISPRi to develop isoform-resolved single-cell Perturb-Seq, a screening approach that enables systematic analysis of alternative promoter function. We hypothesized that alternative promoters are independent regulatory units capable of driving distinct cellular phenotypes and drug responses. By combining promoter-specific targeting with single-cell transcriptional profiling, we uncover widespread functional divergence between alternative promoters of the same gene and demonstrate their potential as therapeutic targets.

## Material and Methods

### Publicly available datasets

We utilised publicly available genome-wide single-cell CRISPR Perturb-Seq data(4) for the cell lines RPE1 and K562. This comprehensive screen utilises over 2 million cells. Each 10x channel (GEM group) was z-normalized to aggregate the whole screen into a gene-by-cell barcode count matrix, in which every cell barcode was associated with a gene knockdown and separated into pseudo bulk populations of cells.

### Deposited Datasets

| Description | Link | Information |
| --- | --- | --- |
| Raw Sequencing data from Perturb-Seq | EBI ArrayExpress<br>E-MTAB-14567 | FASTQ files from CRISPR library and transcriptome |
| Processed data from Perturb-Seq | EBI ArrayExpress<br>E-MTAB-14567 | BAM files output from cellranger |

### Publicly available Datasets

| Cell Line | Description | Identification number | Reference |
| --- | --- | --- | --- |
| RPE1 | Single-cell CRISPR Perturb-Seq data |  | (4) |
| K562 | Single-cell CRISPR Perturb-Seq data |  | (4) |
|  | CRISPRa growth data | Supplementary data from publication | (15) |
| Tissue-wide | Dolcetto sgRNA library |  | (9) |
| Tissue-wide | Fantom 5 CAGE-seq | hg19.cage_peak_phase1and2combined_tpm_ann.osc.txt.gz | (10) |
| K562 | Long Read | ENCFF516IEG | (16) |
| MCF-7 | Tamoxifen | SRP116398 | (17) |
| MCF-7 | Resistance line | SRP262641 | (18) |
| MCF-7 | H3K4me3 ChIP-seq | ENCSR000DWJ GSM945269 | (19) |
| MCF-7 | IDR CAGE-seq | ENCSR000CJO, GSM849364 | (20) |
| MCF-7 | Long-read Pacbio CCS data | See citation | (21) |
| MCF-7 | ESR1 and AR ChIP-seq | GSM1187117 | (22,23) |
| MCF-7 | Tamoxifen treatment | GSE103242 |  |

### CRISPR library and promoter analysis

To identify tissue-wide regulation promoters per a gene, we downloaded RLE normalized CAGE peak expression profiles from Fantom5 (hg19.cage_peak_phase1and2combined_tpm_ann.osc.txt.gz)(10) and used liftover (hg19ToHg38.over.chain.gz) to convert hg19 genome coordinates to hg38. We defined distal alternative promoters as separated by more than 1000 nucleotides. Only promoters contributing more than 20% of total gene expression in at least one tissue type were considered as expressed. CRISPR library sgRNAs within 1000nt of a promoter were considered as targeting the promoter.

To analyse the correlation between knockdown efficiency and several alternative promoters, we downloaded the analysed count data long-read nanopore sequencing data from the ENCODE database from K562(16). We only considered promoters separated by 1000nt, which were considered separate promoters. For promoters within this range, the count data was concatenated.

For the promoter expression analysis, fastq files were obtained for each knockdown from the Repogle et al. RPE1 dataset(4,24) and analysed using the transcript analysis program Whippet. The sgRNA and transcript TPM values concatenated considered all transcripts with promoters within 1000nt of P1 or P2 sgRNA as targeted. All transcripts with promoters outside this range were discarded (for the main figure) or included in additional calculations.

### Analysis of RNA-seq datasets

Whippet was used with an index constructed from Hg38 genome with annotation from Ensembl v102 using default settings. Ensembl v102 annotations supplemented with RefSeq annotations for ESR1. Whippet-quant was run with –biascorrect and default settings. Relative promoter expression was calculated by concatenating transcript TPM for P1 and P2 associated transcripts relative to total gene expression.

### Cloning of Promoter-specific CRISPRi (promCRISPRi) dual guide library for capture

The PromCRISPRi library was generated using a dual guide RNA cloning method, as outlined in Repogle et al. 2020(24) and 2022(4). This method reduces library size and maximizes knockdown using two guides designed to target the same promoter on a single vector. Briefly, oligonucleotides were synthesised in a pool format (Twist Biosciences). SgRNAs protospacer sequences were spaced by a BsmBI enzyme site and flanking sequences of BstX1/BlpI followed by PCR adapters: 5’-PCR adaptor – CCACCTTGTTG – protospacer sequence A – gtttcagagcgagacgtgcctgcaggatacgtctcagaaacatg – protospacer sequence B – GTTTAAGAGCTAAGCTG – PCR adaptor-3’ (Supplementary Table S2).

Oligo pool was PCR-amplified, digested with BstX1/Blp, purified and ligated into pJR85 (Addgene #140095) previously digested with the same enzymes. The newly generated library plasmid was digested with BsmBI enzyme and a second plasmid pJR89 (Addgene #140096) was also digested with the same enzyme to remove CR3/hU6 promoter insert. The released fragment from pJR89 was purified and ligated to the final library plasmid. Final PromCRISPRi library and the guide RNA distribution was confirmed by NGS sequencing(25).

### Cell culture and maintenance

Human embryonic kidney (HEK) 293T, MCF-7 wild-type and MCF-7 dcas9 Krab cell lines were cultivated in Dulbecco′s modified eagle medium (DMEM) supplemented with 10% (v/v) fetal bovine serum (FBS), and 1% penicillin/streptomycin. Cells were maintained at 37°C in a humidified atmosphere containing 5% CO_2_. For all experiments, cells at a lower passage were used, and a mycoplasma test was conducted.

### Lentivirus production and transduction

To produce lentivirus, HEK239T cells, growing in a T-25 flask at 80-90% confluency were co-transfected with 2 µg of pCAG-VSVG, 4 µg of psPAX2 (Addgene #35616, #12260), with either the dCas9 plasmid pHR-SSFV-KRAB-dCas9-P2A-mCherry (Addgene #60954) or the IsoCRISRPi library; using Lipofectamine 3000 transfection reagent (Thermo Fisher Scientific) following the manufacturer’s recommendations. Media was replaced 24 hours after transfection, and 48 hours later, supernatants containing viruses were collected by centrifugation and filtration through a 0.45-μm PVDF filter (Millex Millipore - 0890). The viruses were either used immediately or stored in aliquots at −80 °C.

To generate a stable MCF-7 KRAB dCas9 cell line, MCF-7 wild-type cells in a T-25 flask at 60 - 70% confluency were transduced for 24 hours with lentivirus containing pHR-SSFV-KRAB-dCas9-P2A-mCherry. Polyclonal populations of mCherry-positive cells were sorted using a FACS Aria II Cell Sorter (BD Biosciences). The purity of the recovered populations was >98%, and dCas9 protein expression levels were verified by Western blot (Supplementary Figure 2F).

After reaching 60-70% confluency, stable MCF-7 KRAB dCas9 cells in a T75 flask were transduced with a lentivirus containing the PromCRISPRi library at a low multiplicity of infection (MOI = 0.1) for 24 hours. Subsequently, the cells were washed 10 times with warm PBS1X to eliminate residual viruses, then trypsinized and divided into 2 T75 flasks. The parental CRISPR guide vector pJR85 used for generating the PromCRISPRi brary has two selectable markers – puromycin and blue fluorescent protein (BFP), enabling two rounds of selection to obtain pure populations of cells expressing dCas9 and harbouring the PromCRISPRi library. The first selection was carried out three days post-transduction, the percentage of cells expressing dCas9 (mCherry+) and dual guide RNA (BFP+) was determined, and cells were sorted to near purity using FACS Aria II (BD Biosciences) (Supplementary Figure 2E). Following the sorting of double-positive cells (mCherry+, BFP+), they were pelleted and seeded in a T-25 flask with DMEM, 10% FBS, 1% penicillin/streptomycin, and 1ug/mL puromycin. By the 8th day of drug selection, more than 70% of the MCF-7 KRAB dCas9 population expressed the PromCRISPRi library (BFP+), and a final cell sorting was conducted for single-cell direct capture Perturb-seq.

### Single-cell direct capture Perturb-seq and sequencing

By the 8th day post-transduction, more than 70% of the MCF-7 KRAB dCas9 population expressed the PromCRISPRi library (BFP+), and a final cell sorting (Aria II Cell Sorter - BD Biosciences) was conducted prior single-cell direct capture Perturb-Seq (Supplementary Figure 2E). PromCRISPRi cells with more than 90% viability were prepared as single-cell suspension and loaded into droplet emulsions to two lanes of Chromium Single Cell Chip K, aiming to recover ∼20,000 cells per GEM group = 40,000 in total. Gene expression (GEX) and PromCRISPRi libraries were prepared following 10X Genomics Chromium Single Cell 5’ Kit User Guide v2 (Dual Index) with Feature Barcode technology for CRISPR Screening (CG000510 Rev B). Sequencing was performed on a NovaSeq 6000 (Illumina) according to the 10x Genomics User Guide.

### ESR1 promoter-specific knockdown validation and proliferation assay

The same cloning strategy, lentivirus production, and cell transduction used for the PromCRISPRi library were also applied to ESR1 gene promoter-specific knockdown and non-targeting gRNA controls. The correct sequence of the ESR1 dual guide CRISPRi plasmids was confirmed by Sanger Sequencing. Following transduction, cells expressing gRNAs (+BFP) and dCas9 protein (+mCherry) were FACS sorted as previously described and cultivated under puromycin drug selection. After 6 days, the cells were harvested for total RNA extraction using the RNeasy Mini Plus Kit (74134 - Qiagen) according to the manufacturer’s instructions. cDNA was synthesized using RevertAid RT Reverse Transcription kit (K1691 - Thermo Fisher Scientific) following the manufacturer’s protocol. qRT-PCR was performed using primers for ESR1 P1, P2 (Supplementary Figure 8) GAPDH, and PCR Master Mix Power SYBR Green (4367659 - Thermo Fisher Scientific) on a QuantSudio 7 Flex RT-PCR System (Applied Biosystems). Relative expression levels were calculated using the 2−ΔΔCT method with GAPDH as the endogenous control and non-target control sgRNA dCas9 as a reference sample. This was calculated across two biological replicates and two technical replicates (two cell populations of two dual guide sgRNA-dCas9 targeting P1 or P2) with NTC population for each—significance tested with Wilcoxon Mann-Witney test. qPCR primer sequences are provided in Supplementary Table 4.

To measure the impact of ESR1 P1 and P2 promoter knockdown on cellular proliferation, after FACS sorting cells were seeded at 10% confluency in 96 well plate and cultivated in DMEM supplemented with 10% FBS and 1% penicillin/streptomycin. The media was replaced for the samples treated with 5 mM Tamoxifen, and the drug was added 24 hours after cell seeding. Cell proliferation was assessed using a live-cell Incucyte S3 (Sartorious) imaging system, capturing images at four defined points per well every 4 hours over 5 days. Two independent experiments were conducted with 3 technical replicates (wells) conditions. Analysis was performed using Incucyte S3 Analysis System. Relative phase object confluence is the phase object confluence subtracted from the confluence at 0h. Lmer package was used to fit a second-degree polynomial stats model to each condition and calculated significant difference in confidence intervals (Supplementary Table 5). Doubling time is calculated as below.

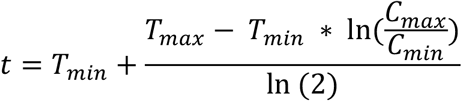

### Promoter identification and guide design

50 candidate genes were targeted in the MCF-7 cell line that were identified as ideal events of alternative promoter usage to be targeted for knockdown via the Perturb-seq technology. There were three stages for library design: 1. Promoter identification, 2. Filtering for the ideal candidates, and 3. Design of guides.

First, the promoter identification through integrating MCF-7 transcriptomic data from IDR of CAGE-seq(20), proActiv(12) of short-read and long-read Pacbio CCS data(21) and annotated with the known promoters refTSS database(26). Next, the top 42 ideal candidates were selected due to being highly expressed in MCF-7 (TPM > 2), containing promoters with >1000 bp distance from each other, RNA transcription factors(27), enrichment of H3K4me3 ChIP-seq(19) and the major promoter upstream of the minor promoter. Finally, guides were designed using FlashFry(28) to the first 100 bp of the promoter of interest. Per promoter (e.g. STAT3 P1), four guides were chosen or, two per dual guide vector.

To estimate a null distribution of expression variability, we used transcriptomic data from cells transduced with non-targeting control sgRNAs (9.5% of the library) to define baseline variability in promoter expression in the absence of targeted perturbation. Non-targeting sgRNAs composed 9.5% of the total library. The guides were targeted at the first 100 bp of a promoter. The top four guides for each site of interest were prioritised if found in Horbleck et al. 2016 hCRISPRiv2.1 guide library, found with a complimentary guide design tool CRISPRDO(29). Finally, they had the highest score of on-target (Moreno-Mateos) and off-target scores (Doench, Table S1 & Figure S2C)(26).

### Gene-level quantification

Using Cell Ranger v8.8.0, the transcriptomic library was aligned to reference GRCh38 v77 using “count”, and the feature library was aligned to the supplied list of guides (Supplementary Table 2). For accurate guide calling, quantification of unique molecular identifier (UMI) into molecule_info.h5 file was reprocessed using guide calling pipeline^8^ for coverage of read per UMI. Two unique sgRNAs expressed from a single lentiviral vector are assigned as ‘ideal’.

### Transcript quantification

Pseudo bulk populations of cells were separated into a specific gene knockdown BAM file. Picard MarkDuplicates removed the PCR-bias read amplification, then BAM files were converted into FASTQ. Whippet was used to quantify transcriptome cDNA with GRCh38. For the transcriptome, transcripts within 800 bp of a promoter were assigned to the promoter and then summed TPM. The percentage of KD calculated between the promoter’s KD of interest and non-targeting control guides. To be defined as a ‘successful KD’, a minimum of 0.5 TPM in both P1 and P2 driven isoforms within the non-targeting control populations was required (n = 31).

### Quantifying transcriptomic differences

Count matrices were filtered for genes with a minimum of 20 cells and genes with a minimum of 1000 reads—highly variable genes assigned to top 2000 with Seurat_v3. From total UMI content normalized, log-scaled expression data, a neighborhood graph was computed (using scanpy.pp.neighbors with n_neighbors=30, method=‘umap’, metric=‘correlation’, and n_pcs=20) followed by UMAP embedding (using scanpy.tl.umap with default parameters). scPerturb(30) to quantify e-statistic using euclidean distance on X_pca. E-test quantified on e-distance to alpha=0.05, runs=200 using the control as the P2 targeted cell population relative to P1. N-terminus change is calculated for every gene if the canonical ATG resides between the P1 and P2 defined co-ordinates (Supplementary Table 1).

Rand score is calculated as how well unsupervised clusters recapitulate P1 and P2 labels. HDBSCAN assigns hierarchical clustering output to identify and label clusters using hyperparameter optimisation. Clusters assigned algorithm=‘best’, cluster_selection_method=leaf, min_cluster_size=10, metric=“manhattan”,alpha=0.6 and allow_single_cluster=False.

### Neighbourhood Gene KD & Differential Gene Expression

Using reference, neighbourhood gene to target gene were identified. Input count matrixes was separated into two pseudo-bulk population between targeted cell population and non-targeting gRNA control, statsmodel z-test with a null hypothesis of no difference in means and standard deviation between the two populations (ddof =1). differentially expressed genes were identified through scanpy rank_genes_group_df t.test method, filtered for genes p-value < 0.05 and log fold change > 0.5.

#### Nanopore Sequencing

MCF7-dCas9 cell lines were transfected with sgRNAs for ESR1-P1, ESR1-P2 and non-targeting control (NTC). The sequencing was undertaken by the Garvan Nanopore Facility using two PromethION flow cells after cDNA conversion and multiplexing. Guppy was used to demultiplex samples.

#### Nanopore Analysis

Nanopore long-read sequencing data was aligned to the hg38 genome using minimap2 (−ax splice). Bambu(31) was used to quantify bam files produced from minimap2 alignment(32) with annotation from refseq database (March 2025, release 229) downloaded from UCSC database. Plotbambu was used to visualize results. differential genes were identified using DESeq2(33) (adj-p-value < 0.05, abs(logFC) > 0.5).

### Spectra Enrichment, FUCCI-cell cycle assignment and CNV score

Spectra(34) was applied to find supervised gene programs without factor analysis biases. The enrichment used 2000 highly variable genes for assignment of gene programs using globally annotated enrichments tested over 1000 epochs, unassigned cell type, lambda = 0.1, delta= 0.001, rho = 0.001 and kappa = None. Cell-cycle assignment was trained using the GSE146773 dataset that pairs cells with known FUCCI-cell states and single-cell transcriptomic data(35). The input count matrix was log-normalised and subsetted for highly variable genes (range of mean 0.0125 to 3 and min. dispersion of 0.5) regulated by the cell cycle, identified previously(36). High-dimensional reduction into UMAP (n PCAs =30). The dimensional reduction is input and split into 0.3 testing and 0.7 training data for classification with kNN scikit-learn model. Hyperparameter optimisation identified best score of 0.8438 (3 s.f.) model (n_neighbours : 11, metric euclidean and uniform weights). ROC Curve used to compare to the scanpy default cell cycle assignment method of score_genes_cell_cycle using known S phase and G2M genes explained previously.

Used inferCNV(37)to detect evidence of chromosomal copy number changes. Used the filtered, normalised matrix across genes (not just highly variable) to compute rolling average gene expression changes for windows of 100 genes with a dynamic threshold of 3 standard deviations for noise filtering (using infercnvpy.tl.infercnv). Genes containing oestrogen or androgen binding sites were identified by using ESR1 and AR ChIP-seq(22,23) overlapping with annotated protein-coding genes.

All figures were plotted using Seaborn and matplotlib in Python or ggpubr on R.

### Survival curve

The pyGEPIA2 package(38) was used to subset 415 Luminal A patients and 192 Luminal B patients from the TCGA/GTEx data source. The hazards ratio is calculated on the Cox PH model with a 95% confidence interval with the group median cutoff for 50% in overall survival from transcripts driven by P1 and P2 (Supplementary Figure 8A).

### Software

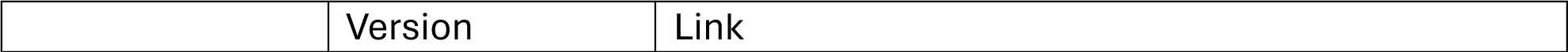
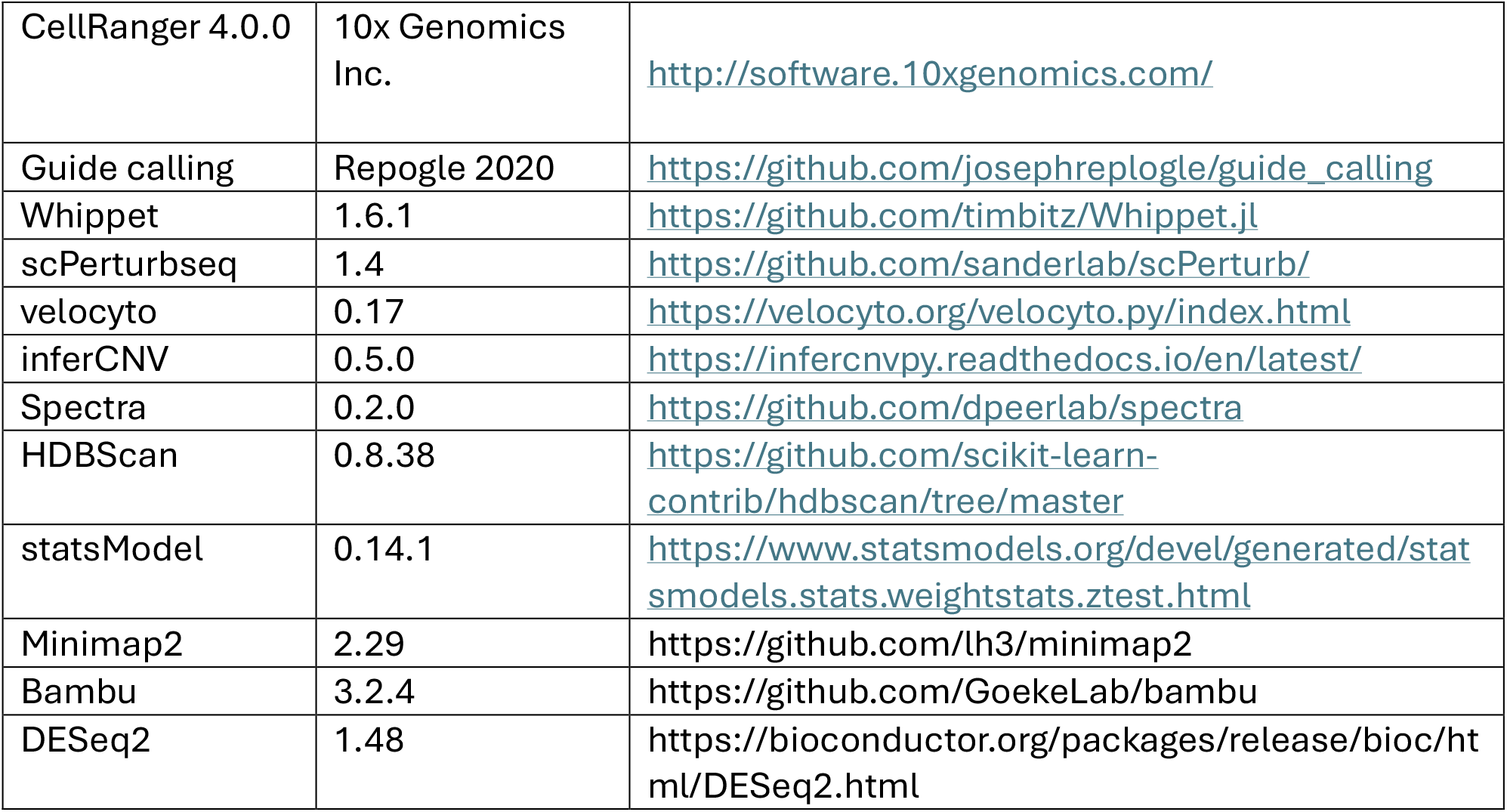

## Results

CRISPRi shows evidence of promoter-specific activity

To understand the scope of alternative promoter regulation in the human genome, we first analysed CAGE-seq data across more than 50 tissue types. This revealed that 14.8% (1,444/9,770) of human protein-coding genes have more than 1000 nucleotides alternative promoters from the major promoter. Given that the CRISPRi machinery has been shown to repress a region of approximately 1000 nucleotides surrounding the sgRNA binding site(14,39), we hypothesized that CRISPRi targeting of these genes might exhibit promoter specificity rather than gene-wide suppression.

To test this hypothesis, we evaluated the relationship between alternative promoter usage and the reported gene knockdown efficiency in published CRISPRi Perturb-Seq datasets from K562 cells(4,24). Using long-read nanopore sequencing data from K562 cells(40) we quantified actively expressed promoters (> 3 supporting reads) for all protein-coding genes. Our analysis revealed a significant inverse correlation between gene knockdown efficiency(4) and the number of alternative promoters expressed per gene (Figure 1A, p < 5.58×10^−37^, Mann-Whitney U-test). This finding suggested that genes with multiple promoters show reduced apparent knockdown efficiency due to continued expression from untargeted alternative promoters.

**FIGURE 1.**
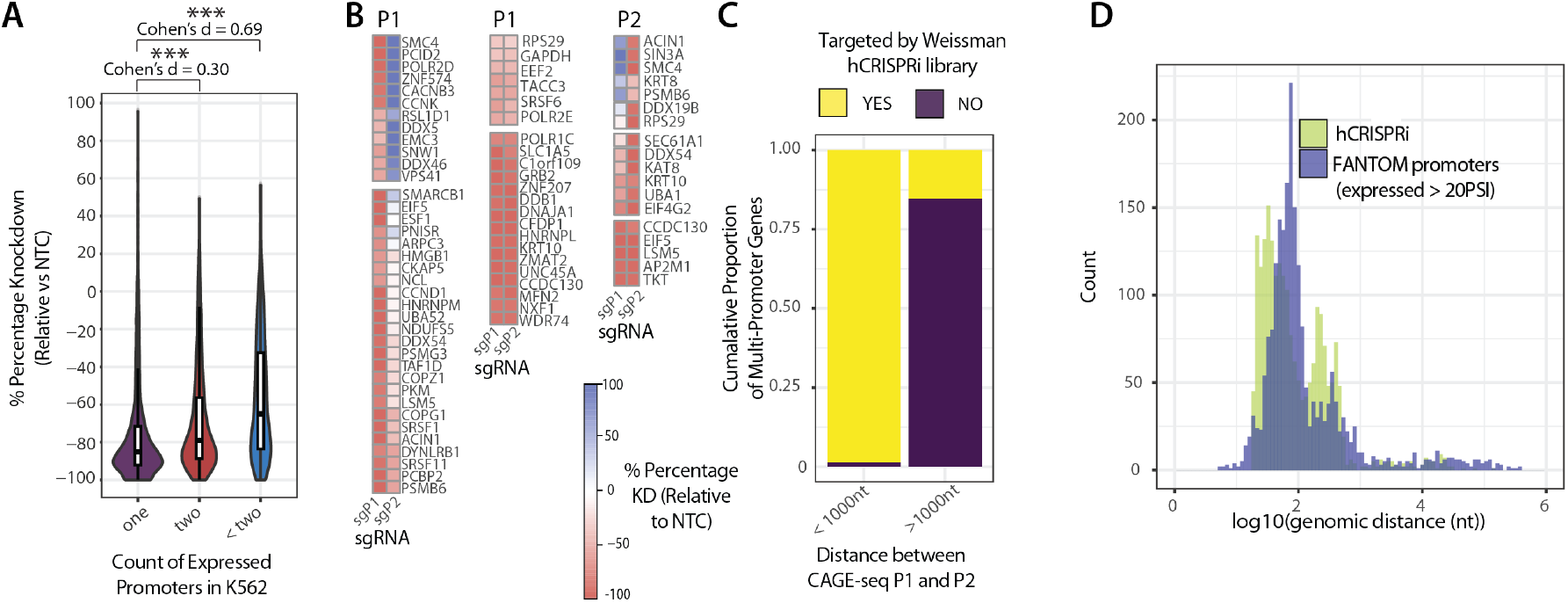
Genome-wide CRISPRi screens overlook alternative promoters. **(A)** Boxplot with outliers removed showing percentage gene knockdown from single-cell CRISPRi-dCas9 Perturb-Seq data, binned based on number of expressed promoters per gene (One vs <two ***P<5.58×10−^37^, One vs two ***P<1.04×10−^15^, Mann–Whitney U-test) across single (n=5,571), dual (n=2,291), and multi (n=778). **(B)** Heatmap showing the relative expression of transcripts initiating from targeted promoters (1 and 2) and non-targeted promoters compared to a negative control from pseudobulk populations. Heatmaps on the left and right display percentage knockdown for sgRNAs targeting P1 and P2 promoters of named genes, respectively. **(C)** Cumulative bar plot showing the fraction of genes targeted by the Weissman sgRNA library, binned into alternative promoters less than and greater than 1000 nt from the major promoter. **(D)** Histogram comparing the maximum genomic distribution of sgRNAs from the Weissman sgRNA library and promoters in a gene body.

### CRISPRi exhibits complex regulatory dynamics at alternative promoters

Detailed examination of transcript-level expression revealed that this reduced efficiency stemmed from two distinct phenomena: persistent expression from untargeted promoters and compensatory upregulation (Figure 1B). While targeted promoters showed consistent suppression, we found that 27.5% of untargeted promoters maintained stable expression (log (FC) < 0.5) compared to normal levels. More strikingly, 11.3% of untargeted promoters displayed increased expression (log (FC) > 0.5), suggesting active compensation for the loss of transcripts from the targeted promoter.

To validate these findings independently, we analysed a CRISPRa dataset, where sgRNAs specifically target individual promoters for activation. Consistent with results from CRISPRi data, we observed an inverse correlation between the number of alternative promoters expressed per gene and the strength of the growth phenotype inhibition (Figure S1A, p < 1.18 × 10^−04^, Mann-Whitney U-test). This parallel finding in both activation and repression contexts is consistent with promoter-specific rather than gene-wide effects of CRISPR-based transcriptional modulation.

### Genome-wide CRISPRi libraries overlook alternative promoters

Given these findings, we evaluated how effectively current genome-wide CRISPRi libraries capture alternative promoter diversity by assessing if promoters identified from CAGE-sequencing data separated by more than 1000 nucleotides are targeted in CRISPRi screens. Analysis of the distribution of sgRNAs from two highly cited genome-wide libraries (Dolcetto(9) and Repogle(4)) revealed systematic gaps in coverage of distal alternative promoters. While these libraries effectively targeted proximal alternative promoters, they failed to target most alternative promoters separated by more than 1000 nucleotides (Figure 1C-D and S1B-C). This includes alternative promoters within 11.5% (n = 1,337) of tested genes that display a tissue-specific switch in major promoter usage (Figure S1D-F). These data suggest that tissue or cell line variability in gene knockdown efficiency may result from alternative promoter usage and that analysis pipelines for CRISPRi/a screens should incorporate alternative promoter usage.

### Isoform-Specific Perturb-Seq enables large-scale transcript-specific analysis

To systematically investigate how alternative promoter usage influences cellular phenotypes, we developed an isoform-specific knockdown screen termed Isoform-Specific Perturb-Seq. We focused our analysis on transcription factors, chromatin modifiers, and RNA binding proteins, reasoning that these regulatory proteins would substantially impact cellular gene expression programs (Figure 2A and S2A). We then selected only genes with promoters more than 1000 nucleotides apart (Figure 2B). Additionally, 9.5% of the library was assigned to non-targeting control sgRNAs, serving as the control comparison and establishing the base rate of promoter expression for the experiment and subsequent analyses.

**FIGURE 2.**
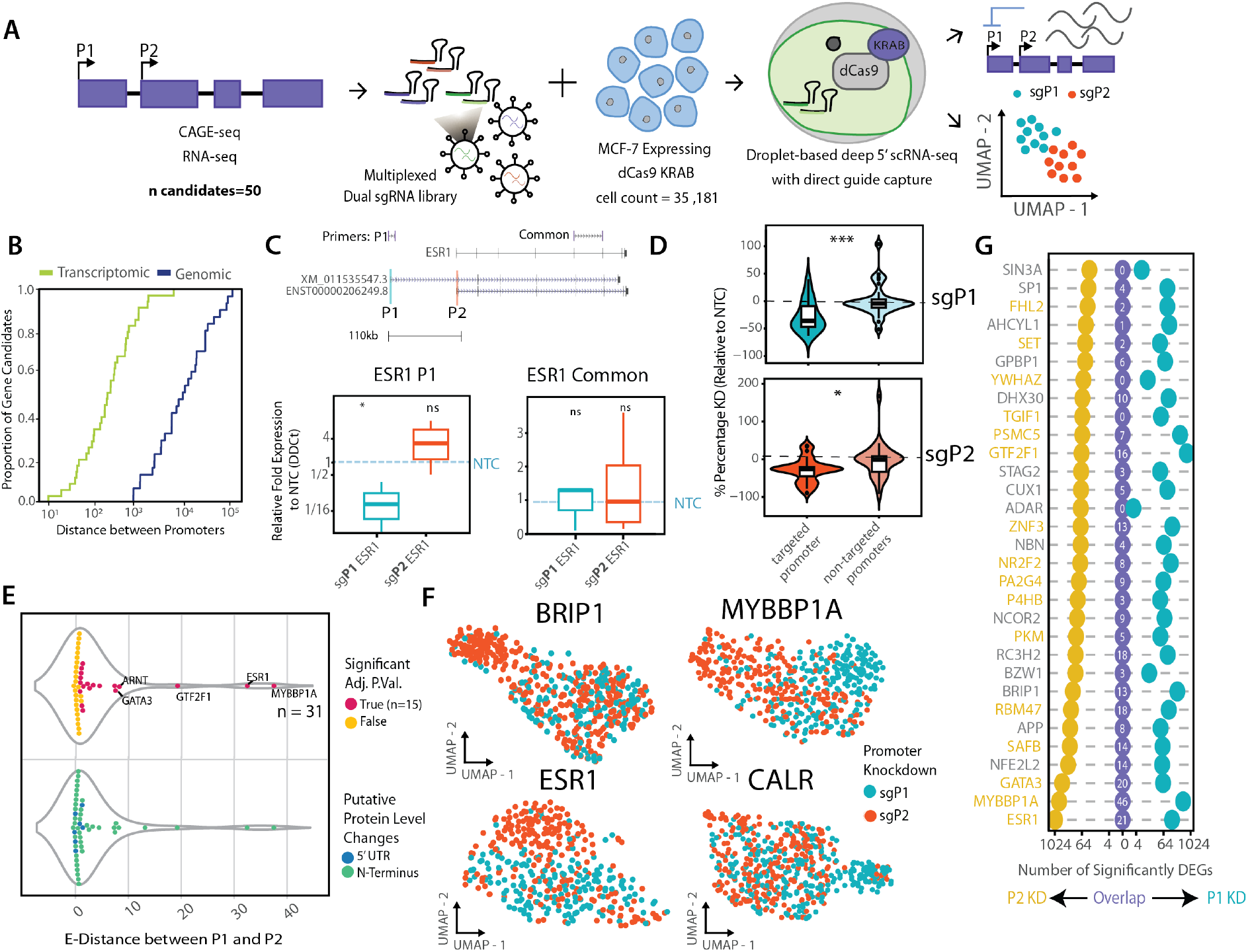
Isoform-Specific Perturb-Seq enables large-scale transcript-specific knockdown. **(A)** A schematic of the pipeline of isoforms-specific Perturb-Seq from promoter identification (left), generation of dual guides for respective P1 and P2 promoters and creation of MCF-7 stable cell line (centre) to 10x single-cell sequencing (right). **(B)** Plot of transcriptome and genomic distances between P1 and P2 promoters for the candidate genes targeted in a screen. The transcriptome distance refers to the distance between the transcription start sites (TSSs) of P1 and P2 in the spliced transcript annotation. **(C)** (TOP) A genome browser track of ESR1 with P1 (green) and P2 (pink) promoters annotated. Top track displaying the targeted primers. Only transcripts with greater than 5 TPM from bulk RNA-seq shown. (BOTTOM) Quantitative RT-qPCR results across sgRNA P1 and P2 from primers targeted to a region common to all estrogen receptor (ESR1) transcripts (LEFT) and primers specific to P1 transcripts (right) pooled from two technical and two biological replicates. Blue line represents the non-targeting control sgRNA relative expression. Relative Expression (ΔΔCt) relative to GADPH, a housekeeping gene (from left to right P=0.0108, P=0.221, P=0.934 and P=0.591, One-sample t-test). The blue dotted line represents the non-targeting control. Significance codes p > 0.05 signified by n.s. (not significant), p<0.05 * **(D)** Violin plots of percentage knockdown of targeted promoters associated transcripts relative to non-targeting controls for sgRNAs targeting P1 promoters relative to transcripts produced from transcripts produced from targeted and non-targeted promoters (n = 31, NTC TPM > 0.5, *** p< 4.7 × 10^−4^, two-sided Welch’s t-test) and P2 promoters (right, *, p< 4.8 × 10^−2^, paired two-sided Welch’s t-test). **(E)** Violin plots showing a statistical quantification of transcriptome distances between single cells with P1-to P2-sgRNAs knockdowns. TOP plot shows adjusted p-values (Monte-Carlo Permutation test with Holm-Sidak Multiple test correction). BOTTOM plot shows putative changes to mRNA or protein transcript isoform defined by if canonical ATG resides between P1 and P2. **(F)** UMAP plots for BRCA1-associated C-terminal helicase 1 (BRIP1), Myb-binding protein 1 (MYBB1A), Estrogen Receptor 1 (ESR1) and Calreticulin (CALR) showing different clustering of single cells knockdown by P1 and P2 promoters. **(G)** A dot plot displaying the differential gene expression (DEG) analysis for P1 and P2 promoters against non-targeting controls. Numbers of P1 and P2 differential genes for each gene are shown with a central number displaying several overlapping significantly differentially expressed genes (genes exp in > 10%, adj. pval < 0.05, log FC > 0.5) between P1 and P2.

To implement our Isoform-Specific Perturb-Seq approach, we generated MCF7 breast cancer cells stably expressing dCas9-CRISPRi. We selected MCF7 cells due to their well-characterized alternative promoter usage and clinical relevance in breast cancer studies. To ensure robust knockdown, we employed a dual guide RNA strategy, targeting each promoter with two sgRNAs (Figure 2A-B, S2, Table S1, Table S2, Methods). Initial validation using quantitative PCR demonstrated successful promoter-specific knockdown of estrogen receptor 1 (*ESR1*), a gene with characterized alternative promoters (Figure 2C, Table S3). We also noted that targeting the P1 promoter of ESR1 significantly reduced the expression of P1-specific transcripts.

Our Isoform-Specific Perturb-Seq screen achieved high-quality single-cell transcriptome data (Median nUMIs per cell = 13,971, Median nGene per cell=3,718) across 27,063 cells (Figure S3-S4). After filtering for genes with significant basal expression in the non-targeting control population (n =31, TPM > 0.5), we observed a significant reduction in transcripts associated with targeted promoters while maintaining expression from untargeted promoters (P1: p < 4.7×10^−4^ ; P2: p < 4.6×10^−2^, paired two-sided Welch’s t-test, Figure 2D, S5A-B and Table S4). This validated our ability to target individual promoters across a large-scale single-cell screen.

### Alternative Promoters drive distinct gene expression programs

Analysis of single-cell transcriptomes revealed that alternative promoters within the same gene can drive subtly different expression programs. Due to the sparsity and zero-inflation characteristic of single-cell RNA-seq data, volcano plots are not well suited to summarize differential gene expression at single-cell resolution. Instead, we applied dimensionality reduction and e-distance metrics that better capture global transcriptomic divergence between promoter knockdowns. After removing quality controls removed lowly expressed genes and transcripts, we used e-distance metrics to quantify transcriptional divergence(30), and found that 48.3% (15/31) of genes with alternative promoters showed significant differences, albeit with small effect size, in cellular transcriptomes when comparing P1 versus P2 knockdown (Figure 2E), with 18% (6/31) showing stronger effects on transcriptome and the rest exhibiting more subtle changes (Figure 2E). These differences showed strong reproducibility between replicates (Figure S5C, Table S5 & Table S6) and compared to non-targeting control transcriptomes (Figure S6), confirming their biological significance.

Analysis of screen-positive genes revealed that the most functionally impactful promoter knockdowns involved transcripts contributing at least 5% of the total gene expression in MCF-7 cells. Notable examples include *ESR1* and Myb-binding protein 1A (*MYBBP1A)*, where knockdown of P1 versus P2 promoters produced distinct transcriptional landscapes (Figure 2F), suggesting that these promoters drive distinct gene expression programs within the total expression landscape of each gene. differential gene expression analysis revealed minimal overlap between genes affected by P1 versus P2 knockdown, suggesting these promoters regulate distinct downstream pathways (Figure 2G). This functional divergence between promoters of the same gene highlights a previously underappreciated layer of transcriptional regulation.

To orthogonally validate the promoter-specific knockdown, we undertook nanopore long-read sequencing of MCF-7 cells transduced with sgRNAs targeting P1 and P2 promoters, as well as a non-targeting control. This analysis validated the promoter-specific knockdown on CRISPRi on transcripts initiated by the respective promoters (Figure S6F). Furthermore, differential gene expression validated the minimal overlap of genes regulated by P1 versus P2 knockdown (Figure S6G) further supporting that promoters often drive distinct downstream pathways.

### Alternative Promoters regulate distinct cellular phenotypes

To understand the biological significance of promoter-specific regulation, we employed Spectra(34), to identify coordinated gene expression programs associated with each promoter. Spectra analyzes single-cell RNA-seq data to calculate pathway activity scores for individual cells, enabling comparison of pathway activation across different promoter knockdowns. These cell-level pathway scores are then statistically evaluated to identify differentially activated biological processes revealing promoter knockdown specific pathway alterations. This approach revealed that alternative promoters can control distinct biological processes, particularly in cell cycle regulation and proliferation pathways (Figure 3A, Table S7).

**FIGURE 3.**
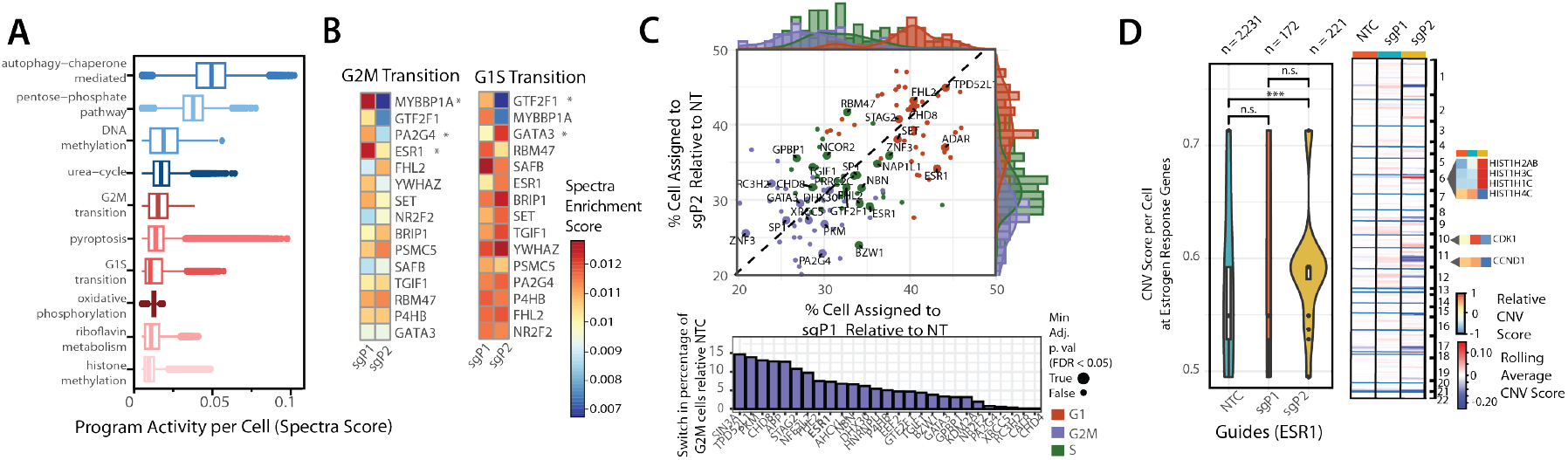
Alternative promoters drive distinct functional gene expression programs. **(A)** A boxplot of the top enriched functional programs (y-axis, *n* = 10) identified by program activity per cell (Spectra score) across the screen. Higher program activity indicates a stronger presence or activation level of a specific gene program within a cell, with values > 0.05 generally signifying a ‘positive’ (active) state for that program. **(B)** A heatmap of the mean Spectra enrichment score for target gene candidates (*n* = 15; successful knockdown and significantly different e-distance scores) comparing P1- and P2-promoter-targeted cell populations across G2M (left) and G1S (right) transitions (two-sided Welch’s t-test, *p* < 0.05). **(C)** (TOP) A scatterplot displaying the percentage of cells in each cell cycle stage (G1 = purple, S = red, G2M = green) for genes targeted in the screen. A dashed line is fitted to negative control guides (two-sided Welch’s t-test, Benjamini/Hochberg FDR). (BOTTOM) A barplot showing the minimum percentage change in G2M occupancy, relative to non-targeting control (NTC), for genes with promoters showing differential effects. *ESR1* is highlighted in bold. **(D)** (LEFT) Violin plots with overlaid boxplots showing putative copy number variants inferred from *infercnvpy* for pseudo-bulk populations of *ESR1* sgP1, sgP2, and NTC. Statistical comparisons: NTC vs sgP2, \*\**P* = 1.1 × 10−^7^; NTC vs sgP1, n.s. *P* = 8.3 × 10−^1^; sgP1 vs sgP2, n.s. *P* = 5.5 × 10−^3^ (Mann–Whitney U-tes*t*). (RIGHT) Heatmap visualization of copy number variants per chromosome, highlighting individual genes by Relative CNV score and Rolling Average CNV score.

Cell cycle phase was assigned using a random forest classifier trained on the highly variable gene (n =1000) expression profiles from cells with known FUCCI-based phase labels (GSE146773) (35,41). This trained model was then applied to the dataset to predict the cell cycle phase for each cell (Figure S7). This analysis demonstrated that 34.38% of alternative promoters significantly altered cell cycle state distributions compared to controls (Figure 3B, p < 0.05, two-sided Welch’s t-test, Benjamini/Hochberg FDR). The most dramatic effects were observed with ESR1, where P1 and P2 promoter knockdowns produced opposing effects on cell cycle progression (Figure 3B, A and Table S8). This suggests that the knockdown of alternative promoters can lead to differences in cell cycle progression.

These transcriptional differences manifested in measurable phenotypic changes. Using inferCNV analysis (*https://github.com/broadinstitute/inferCNV*), which is computational tool used to analyze single-cell RNA sequencing (scRNA-seq) data to infer large-scale chromosomal copy number variations (CNVs), we identified significant increases in copy number variations among cell cycle-related genes specifically in the P2 knockdown population, compared to P1 and control (Figure 3D, Table S9 and S8D, p < 1.1×10^−7^, Wilcoxon rank sum test) with CNVs identified in multiple cell cycle related genes (Figure 3D – right). Live cell imaging confirmed these findings, showing that P2 knockdown significantly reduced proliferation rates ((Figure 3B, p < 1.62 × 10^−10^, t=-6.614, linear-mixed model, Table S10-11), while P1 knockdown had minimal effect on cell growth (Figure 3B, p < 0.0798, t=1.757, linear-mixed model, Table S10-S11). These findings suggest that the expression of the two *ESR1* promoters exert divergent effects on cellular proliferation (Figure 3B and S9A).

### ESR1 promoter specificity determines patient outcomes

Given the clinical importance of estrogen receptor signalling in breast cancer(42), we investigated the relationship between ESR1 promoter usage and patient outcomes. Analysis of nanopore long read RNA-seq data(43,44) across a range of MCF-7 cells revealed coordinated splicing between the P1 promoter and an alternative last exon(45) (Figure S8E). This results in the P1 and P2 promoters driving the production of distinct protein isoforms with coding differences in the activation function-2 (AF2) domain (Figure 4A), a region responsible for binding estrogen, transcriptional coactivators, and selective estrogen receptor modulators (SERMs)(46). The P1 isoform is predicted to contain a disordered C-terminal domain that potentially modifies interactions with estrogen and SERMs, while the P2 isoform produces the canonical ESR1 protein.

**FIGURE 4.**
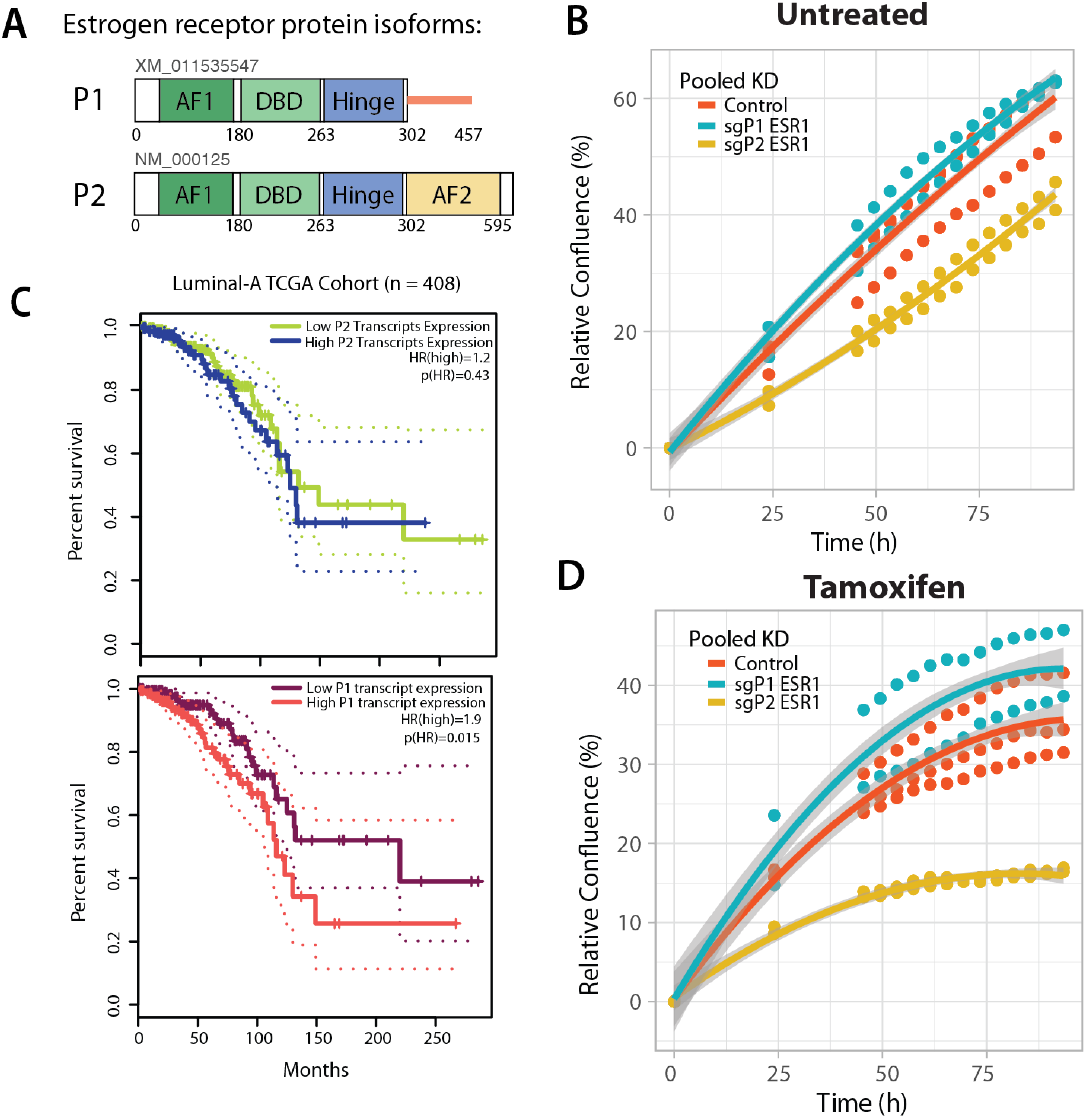
Promoters of the estrogen receptor display tamoxifen-specific responses. **(A)** Domain structures of the two major transcript isoforms of ESR1 based on RNA-seq data AF1 = activation function-1 domain; DBD = DNA-binding domain; AF2 = activation function-2 domain. **(B)** Cellular proliferation results using Incucyte for control-, P1- and P2 knockdown cells untreated (linear mixed model, sgP1 v Control p < 0.0798, sgP2 v Control n.s. p <1.62 × 10^−10^, grey showing confidence interval (CI) Table S11). **(C)** Kaplan-Meier (KM) plots displaying survival curves for patients with Luminal-A breast cancer. KM plots showing transcripts originating from ESR1 promoter 1 (P1) on the left and ESR1 promoter 2 (P2) on the right. Showing low (<median) and high (>median) transcript expression (Hazard Ratio, Cox PH Model). **(D)** Line plot showing cellular proliferation results in MCF-7 breast cancer cells upon control-, P1- and P2-knockdown lines upon tamoxifen treatment (5µM) (linear mixed model, sgP1 v Control p < 1.86 × 10^−5^, sgP2 v Control p < 2.00 × 10^−16^, grey showing confidence interval (CI) Table S11).

Analysis of TCGA data from 299 breast cancer patients(47) revealed a striking promoter-specific relationship with survival. High expression from the P1 promoter strongly correlated with decreased survival in luminal-A breast cancer patients (HR = 1.9, p < 0.015, Figure 4C), while P2 promoter expression showed no significant association (HR = 1.2, p = 0.43) (Figure 4C). This relationship was specific to the luminal-A subtype(48), as no significant correlation was observed in luminal-B breast cancer (Figure S9E), indicating a potential subtype-specific association for ESR1 promoter usage.

### differential drug response of ESR1 promoter isoforms

To understand the mechanistic basis for these clinical observations, we examined ESR1 promoter usage in the context of drug response. RNA-seq data analysis from tamoxifen-resistant MCF7 cells(18) revealed a significant activation of P1 promoter usage compared to parental cells (Figure S9B-C). Resistant cells showed a 2.3-fold increase in P1 promoter usage (p < 0.01) with a corresponding decrease in P2 promoter activity (Figure S9B-C). This suggests that cells may become resistant to tamoxifen by altering their ESR1 promoter usage.

To directly test the functional impact of promoter-specific expression on drug response, we performed tamoxifen sensitivity assays in cells with P1 or P2 knockdown. Strikingly, P1 and P2 knockdown produced opposing effects on tamoxifen sensitivity. P2 knockdown cells showed heightened sensitivity to tamoxifen, with a significant reduction in proliferation rate compared to control cells (Figure 4D, p < 2.00 × 10^−16^, t=-10.998 linear-mixed model, Table S10). In contrast, P1 knockdown cells exhibited increased proliferation in the presence of tamoxifen, growing significantly faster than control cells (Figure 4D, p < 1.86 × 10^−5^, t=4.38, linear-mixed model, Table S10). These divergent responses were specific to tamoxifen treatment, as baseline proliferation rates showed different patterns (Figure 3C), indicating a potential association between this promoter and isoform expression in drug resistance.

Notably, the P1 promoter shows tissue-specific expression patterns (Figure S9D), with high activity in breast tissue. This tissue specificity, combined with its selective role in drug resistance, indicates that targeting P1-specific transcripts could provide a more precise therapeutic strategy than current approaches that broadly target all ESR1 isoforms.

## Discussion

### Limitations of Genome-Wide Functional Screens and the Importance of Promoter-Level Analysis

Our findings advance our understanding of transcriptional regulation by systematically demonstrating that alternative promoters within the same gene can drive distinct cellular programs. This work exposes a critical limitation in current CRISPR-dCas9 functional screens(14,39), where the regulatory complexity and potential compensatory mechanisms between alternative promoters are often overlooked. The discovery that 47% of surveyed transcription and chromatin factors drive distinct transcriptional programs suggests that current screening approaches may miss important regulatory dynamics, particularly in the context of gene regulation. An important note, though, is that differential expression in single-cell data must be interpreted cautiously due to the inherent sparsity, variability, and lack of true biological replicates. While we adopted conservative thresholds consistent with field standards, small fold changes may reflect meaningful shifts in regulatory programs rather than strong individual promoter effects. Our use of a large non-targeting control population, empirical null distributions, and phenotypic validation supports the robustness of our conclusions.

Moreover, we demonstrate that untargeted promoters can exhibit compensatory upregulation, maintaining overall gene expression despite successfully targeting individual promoters. This compensation mechanism suggests that traditional measures of knockdown efficiency may underestimate the effectiveness of CRISPR-based perturbations when applied to genes with multiple promoters. These findings emphasize the need to incorporate promoter-level analysis into the design and interpretation of functional genomics studies.

### Role of Alternative Promoters as Drivers of Unique Cellular Phenotypes

Our isoform-specific Perturb-Seq approach reveals a remarkable autonomy of alternative promoters in driving distinct cellular phenotypes and is complementary to recent efforts to use engineered zinc fingers to drive differential activation of alternative promoters(49). This autonomy enables cells to generate diverse transcriptional landscapes without altering their genomic sequence, a feature particularly evident in our analysis of ESR1, where distinct promoters showed opposing effects on cell cycle progression and drug response. The ability of alternative promoters to integrate diverse regulatory signals - from transcription factors to chromatin modifications - positions them as sophisticated regulatory nodes that fine-tune cellular responses to environmental cues(11,12,50). This functional divergence suggests that promoter selection represents a fundamental mechanism for generating cellular diversity rather than mere transcriptional redundancy.

### Therapeutic Opportunities of Targeting Alternative Promoters in Cancer

The dysregulation of alternative promoter usage is emerging as a critical mechanism in cancer progression and therapeutic resistance(12,51). The clinical relevance of promoter-specific regulation is particularly evident in our analysis of breast cancer, where we discovered that ESR1 promoters differentially impact tamoxifen sensitivity, with P1 knockdown conferring resistance while P2 knockdown enhanced drug sensitivity. This finding aligns with clinical data showing that P1 overexpression correlates with decreased survival in luminal-A breast cancer patients, suggesting that promoter usage patterns could serve as predictive biomarkers for treatment outcomes. Although P1-driven transcripts represent a substantial proportion of total ESR1 expression in tamoxifen-resistant cells, these data alone do not establish that P1 expression is functionally causal in resistance. The observed association may reflect broader transcriptional reprogramming or serve as a surrogate marker for other upstream regulatory changes. Functional studies beyond transcript-level perturbation and cellular proliferation assays are required to determine the phenotypic relevance of P1 expression in this context. Nevertheless these results suggests that, more generally, monitoring promoter usage patterns could help stratify patients for targeted therapies

Our findings also open other therapeutic possibilities. For example, our analysis revealed that the P1 promoter exhibits tissue-specific expression patterns suggesting selective targeting could reduce systemic side effects commonly associated with current estrogen receptor-targeted therapies(42). This expands on the idea that with isoform-specific targeting, we may achieve more precise therapeutic control while minimizing off-target effects compared to broadly inhibiting gene function(52).

Moreover, our observation that specific promoters can drive drug resistance suggests new therapeutic strategies. For example, in breast cancer, targeting specific ESR1 promoters in combination with standard endocrine therapy might improve treatment outcomes by preventing the emergence of resistance through promoter switching(46,53,54). Understanding compensatory promoter activation could also inform strategies to prevent or overcome drug resistance, potentially through combination therapies that target both primary and compensatory mechanisms(46,53,54).

### Future Directions and Technological Implications

Our work establishes a framework for systematic analysis of alternative promoter function, but several important questions remain unexplored. Future research should examine the tissue-specific nature of alternative promoter usage and its implications for drug development. Understanding the mechanisms governing compensatory promoter activation and developing improved CRISPR libraries that comprehensively target alternative promoters will be crucial. Additionally, investigating how alternative promoter usage changes during disease progression and treatment could provide valuable insights for therapeutic development. An important limitation of our work is that we did not impose a minimum expression cutoff during guide design, in hindsight, applying a threshold—such as requiring promoters to contribute at least 5% of total gene expression—may have been a valuable refinement, as the most phenotypically active promoters in our screen exceeded this level.

The integration of promoter-level analysis into drug discovery pipelines could significantly enhance our ability to develop more effective and precise therapeutic strategies. This is particularly relevant for diseases where isoform diversity plays a crucial role, including cancer, neurodegenerative disorders, and autoimmune conditions(55-57).

## Conclusion

In conclusion, our Isoform-Specific Perturb-Seq approach reveals a critical yet underappreciated role of alternative promoters in generating cellular diversity. This suggests that targeting a single promoter per gene may obscure critical regulatory dynamics or yield biologically misleading outcomes. This insight implies that certain dCas9 screens may have missed important regulatory elements, potentially overlooking the complexity of gene expression regulation. Our results also highlight the potential of leveraging dCas9 technology to develop isoform-specific therapies(52), enabling precise modulation of individual protein isoform expression thus paving the way for more targeted and effective therapeutic strategies.

## Acknowledgements

We acknowledge the Garvan-Weizmann Centre for Cellular Genomics team for their assistance with the single-cell assay. Special thanks to Chia-Ling Chan, Winnie Luu, Yasmin Husaini, Eric Lam, Hanjie Wubvand, and Dominik Kaczorowski. We also acknowledge the UNSW Restech HPC Scheme DOI: 10.26190/PMN5-7J50 for computational support. We are grateful to all members of the Weatheritt lab past and present for their critical assistance with this paper and all their support.

## Author contributions

Conceptualization (Robert Weatheritt), Resource (Robert Weatheritt), Data Curation (Helen King, Tim Sterne-Weiler and Robert Weatheritt), Software (Helen King), Formal Analysis (Helen King, Daisy Kavanagh, Savannah O’Connell, Tim Sterne-Weiler and Robert Weatheritt), Supervision (Helaine Graziele S. Vieira, Timothy Sterne-Weiler, and Robert J Weatheritt), Funding acquisition (Robert Weatheritt), Validation (Helen King, Savannah O’Connell, Daisy Kavanagh, Sofia Mason, Cerys McCool, Javier Fernandez-Chamorro, Christine Chaffer, Susan J Clark, Helaine Graziele S. Vieira, Timothy Sterne-Weiler, and Robert J Weatheritt), Investigation (Helen King, Helaine Graziele S. Vieira, Timothy Sterne-Weiler, and Robert J Weatheritt), Visualization (Helen King, Robert Weatheritt), Writing – original draft (Robert Weatheritt), Project administration (Helaine Graziele S. Vieira and Robert Weatheritt), Writing – review & editing (all authors).

